# Distinct phospho-variants of STAT3 regulate naïve pluripotency and developmental pace *in vivo*

**DOI:** 10.1101/2022.03.08.483469

**Authors:** Takuya Azami, Sophie Kraunsoe, Graziano Martello, Yihan Pei, Thorsten Boroviak, Jennifer Nichols

## Abstract

STAT3 has been studied extensively in the context of self-renewal of naïve pluripotent mouse embryonic stem cells. We uncovered acute roles for STAT3 and its target, TFCP2L1, in maintenance of epiblast and primitive endoderm during *in vivo* diapause. On an outbred genetic background, we observed consistent developmental retardation from implantation until embryonic day 11.5, beginning with significant reduction of epiblast cells at implantation in *Stat3* null embryos. Remarkably, mutants closely resemble non-affected embryos from the previous day at all postimplantation stages examined. We attribute this phenotype to loss of the active serine phosphorylated form of STAT3 required for neural differentiation and implicated in growth defects in mice and humans. Bulk RNA-sequencing analysis of isolated epiblasts revealed compromised lipid metabolism in *Stat3* null embryos by embryonic day 6.5. Furthermore, we demonstrate that gastruloids generated from *Stat3* null ESCs failed to extend the posterior axis or maintain BRACHYURY expression and were underrepresented in this region when mixed with wild type cells in chimaeric gastruloids. Our study implicates a role for STAT3 in temporal control of embryonic progression and metabolic mechanisms.

## Introduction

The naïve pluripotent epiblast, founder of the foetus, is established in the murine embryo by the time of implantation and becomes ‘primed’ for germ layer segregation as it subsequently expands (Nichols and Smith, 2009). These two phases of pluripotency can be captured *in vitro* as self-renewing stem cell lines (Brons et al., 2007; Evans and Kaufman, 1981; Martin, 1981; Tesar et al., 2007). Embryonic stem cells (ESCs), derived directly from preimplantation murine embryos, can be maintained indefinitely in an undifferentiated, ‘naïve’ pluripotent state in medium supplemented with leukaemia inhibitory factor (LIF) (Smith et al., 1988; Williams et al., 1988), operating via signal transducer and activator of transcription (STAT)3 (Burdon et al., 1999; Matsuda et al., 1999; Niwa et al., 1998). The LIF receptor (LIFR) forms a complex with glycoprotein (gp)130, also known as IL6ST, and activates Janus-associated kinases (JAKs). JAKs phosphorylate STAT3 in its cytoplasmic monomeric state at tyrosine 705 (pY705) (Ni et al., 2004; Zhang et al., 2000). In addition, STAT3 can be phosphorylated at serine 727 (pS727) by mitogen-activated protein kinases (MAPK) (Huang et al., 2014). Mutation of pY705 ablates ESC self-renewal in standard (serum/LIF) culture conditions, whereas pS727 is not required prior to neuronal differentiation (Huang et al., 2014). Recent studies implicate phosphorylation of Y705 for importation of STAT3 into mitochondria, while S727 phosphorylation is needed for mitochondrial STAT3 transcriptional activity (Carbognin et al., 2016; Gough et al., 2013; Peron et al., 2021). STAT3 mutations are associated with postimplantation developmental defects in mouse and human (Gutierrez, 2020; Shen et al., 2004).

LIF or related cytokines were previously considered essential for ESC self-renewal. However, a refined culture regime, known as ‘2i’, based upon dual inhibition of Glycogen Synthase Kinase (GSK)3 and MEK/ERK, obviates requirement for STAT3 signalling. ESCs deficient for STAT3 could be derived from mutant embryos by incubation from the morula stage in these inhibitors (Ying et al., 2008). Nevertheless, addition of LIF enhances efficiency of wild type ESC propagation, particularly during expansion from single cells (Nichols and Boroviak, 2016; Nichols et al., 2009; Wray et al., 2010). *Stat3* null ESCs enabled analysis of downstream signalling events associated with ESC self-renewal, and thus characterisation of the signalling network operative during maintenance of naïve pluripotency *in vitro* (Martello et al., 2013; Ye et al., 2013). The most significant player emerging from this analysis was TFCP2L1 (also known as CRTR-1) whose forced expression could rescue *Stat3* null ESCs in serum/LIF culture. Using pathway analysis and computational modelling an essential role for TFCP2L1 in ESC maintenance was proposed and supported experimentally (Dunn et al., 2014). Moreover, transfection of *Tfcp2l1* into epiblast stem cells (EpiSCs) derived from postimplantation epiblasts (Brons et al., 2007; Tesar et al., 2007) could direct reprogramming from primed to naïve pluripotency, confirming participation of TFCP2L1 in the naïve pluripotency network (Martello et al., 2013; Ye et al., 2013). However, its function for self-renewal *in vivo* or *de novo* ESC derivation was not explored.

Combined maternal and zygotic deletion revealed an essential requirement for STAT3 during blastocyst expansion, confirming its suspected function in epiblast formation (Do et al., 2013). Perdurance of maternal STAT3 protein in zygotic *Stat3* null embryos, however, permits developmental progression beyond cleavage stages. These embryos could implant in the uterus, but rapidly acquired abnormalities (Takeda et al., 1997). In normal laboratory rodents, the state of naïve pluripotency is relatively transient, lasting no more than two days. It is therefore debatable whether self-renewal of stem cells occurs at this stage during uninterrupted development. Murine preimplantation embryogenesis can be prolonged by diapause, a natural, facultative phenomenon occurring when a dam conceives whilst suckling a previous litter. Embryos progress to the peri-implantation stage, but delay implantation until a source of oestrogen is regained. Diapause can be achieved experimentally by ovariectomy prior to the physiological burst of oestrogen secretion at embryonic day (E)2.5 (Weitlauf and Greenwald, 1968).

We report an obligation for STAT3 and TFCP2L1 in epiblast maintenance during diapause. Furthermore, by crossing *Stat3* heterozygous mice from the original genetic background (Takeda et al., 1997) to the outbred CD1 strain, we extended postimplantation development of *Stat3* null embryos beyond gastrulation and thereby uncovered a striking role for STAT3 in temporal control of embryonic progression and metabolic mechanisms.

## Results

### STAT3 pY705 and TFCP2L1 peak in epiblast during embryonic diapause

We confirmed distribution of pY705 in preimplantation embryos (Do et al., 2013) by immunofluorescence (IF), showing nuclear localisation from 2-cell to late blastocyst stages (Fig. 1A). Whereas the pluripotency-associated factor NANOG is highly expressed exclusively in the epiblast at E4.5 (Silva et al., 2009) levels of pY705 and TFCP2L1 were visibly diminished (Fig. 1A). The potential roles of STAT3 and TFCP2L1 in sustaining pluripotency *in vivo* were investigated via induction of diapause, as described previously (Nichols et al., 2001). Embryos were recovered 4 days after ovariectomy, performed at E2.5. Nuclear levels of pY705 and TFCP2L1 increased dramatically in epiblast (Fig. 1B), consistent with the transcriptional resemblance of diapause epiblast to self-renewing ESCs *in vitro* (Boroviak et al., 2015). We developed an image analysis tool to enable robust quantification of pY705 and TFCP2L1 protein distribution in peri-implantation and diapause embryos (Fig. 1C,D; Fig. S1A) and thereby confirmed that both proteins become enriched in ICM cells during diapause (Fig. 1E,F; Fig. S1B,C).

**Figure 1.**
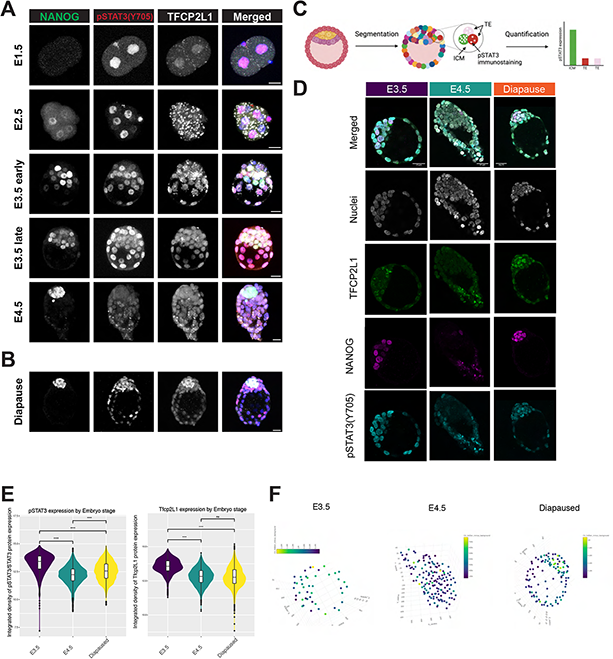
Distribution of pSTAT3(Y705) and TFCP2L1 in preimplantation and diapause embryos. (A, B) Confocal images of NANOG, pSTAT3(Y705), and TFCP2L1 immunofluorescence in preimplantation (A) and diapause embryos (B). Scale bar = 20 μm. (C) Schematic of image analysis pipeline to segment individual nuclei in 3D and quantify fluorescence of each channel. (D) Immunofluorescence staining of E3.5, E4.5 and diapause embryos for DAPI, TFCP2L1, NANOG and pSTAT3(Y705). Scale bar = 25 μm. (E) Violin plots with overlaid boxplots of the integrated density of pSTAT3(Y705) and TFCP2L1 expression across each nucleus from each embryo by stage (E3.5, n = 29, E4.5, n= 33, Diapause, n = 54). Segmented nuclei were filtered for DAPI signal and volume to remove any erroneous segmentation. * *P* < 0.01. (F) Representative example embryos from each developmental stage with nuclei represented by a scatter point in Fig. S1B and colour coded based on pSTAT3(Y705) integrated density.

### Propagation of naïve pluripotency *in vivo* requires STAT3 and TFCP2L1

To assess functionality of STAT3 signalling during maintenance of naïve pluripotency, heterozygous mice were mated *inter se* to generate wild type (WT), heterozygous (het) and mutant (null) embryos for *Stat3*. Embryos were recovered at E4.5, and IF for NANOG, pY705 and TFCP2L1 performed. NANOG and TFCP2L1, but no pY705 STAT3 protein, were detected in epiblasts of null embryos (Fig. 2A). Whereas WT and het embryos retrieved 4 days after ovariectomy possessed large ICMs with many NANOG-positive cells, embryos lacking STAT3 exhibited complete loss or severe reduction of the whole ICM (Fig. 2B,C). Similarly, late blastocysts lacking *Tfcp2l1* exhibited NANOG and pY705 IF (Fig. 2D), but diapaused null embryos had no discernible ICM (Fig. 2E,F). *Tfcp2l1* null embryos were underrepresented, which may be attributed to loss before or during retrieval, or genotyping errors of the parents. Taken together, these results implicate immediate requirement for STAT3 signalling during diapause.

**Figure 2.**
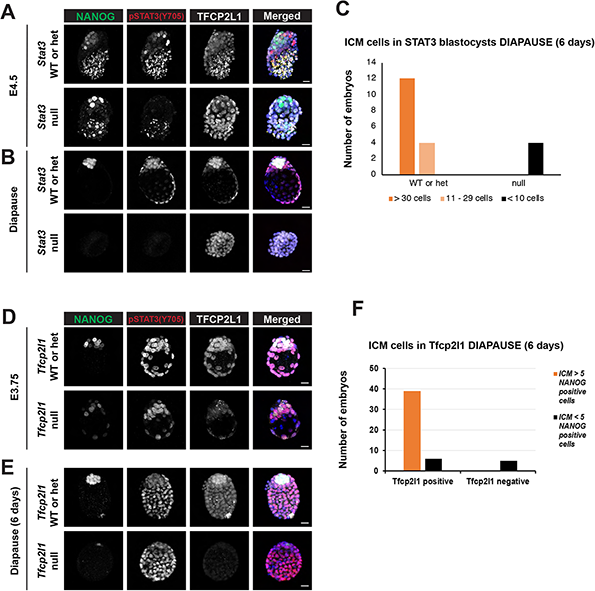
STAT3 and TFCP2L1 are required for ICM maintenance during diapause. (A, B) Immunofluorescence of NANOG, pSTAT3(Y705), and TFCP2L1 in E4.5 (A) and diapause (B) *Stat3* WT/het and null embryos. Scale bar = 20 μm. (C) Number of ICM cells in *Stat3* WT/het and null diapause embryos in (B). (D, E) Immunofluorescence of NANOG, pSTAT3(Y705), and TFCP2L1 in E3.75 (D) and diapause (E) *Tfcp2l1* WT/het and null embryos. Scale bar = 20 μm. (F) Number of ICM cells in *Tfcp2l1* WT/het and null diapause embryos in (E).

Our previous derivation protocol (Ying et al., 2008) was used to capture CD1 background ESCs. Nine WT, 32 het and 13 null ESC lines were generated from 56 embryos by *inter se* mating of *Stat3* het mice (Table S1). Each embryo was genotyped by PCR using lysed trophectoderm produced during immunosurgery (Nichols et al., 1998; Solter and Knowles, 1975). Conversely, from 65 morulae generated from *Tfcp2l1* heterozygous intercross, 6 WT and 49 het, but no null ESC lines were derived (Table S1), suggesting a distinct requirement for TFCP2L1 in capture of pluripotency *in vitro*.

### STAT3 is dispensable for organogenesis but required to sustain developmental pace

Significantly fewer NANOG-positive cells were observed in *Stat3* null embryos at E4.5, whereas GATA6 cell numbers did not differ appreciably between genotypes (Fig. 3A,B). However, we noted dramatic reduction in the mean number of trophectoderm cells (counted by position in the embryo and absence of NANOG and GATA6) in mutant embryos compared with WT or hets (Fig. 3B), which we attribute to decreased FGF signalling resulting from the smaller size of null epiblasts; we previously demonstrated requirement for FGF in trophectoderm proliferation (Nichols et al., 2009; Nichols et al., 1998). Dissection of embryos from *Stat3* het inter-cross mating after implantation revealed persistence of developmental delay in *Stat3* nulls equivalent to around one day compared with their littermates until at least E11.5, with no other obvious morphological defects (Fig. 3C; Table S2).

**Figure 3.**
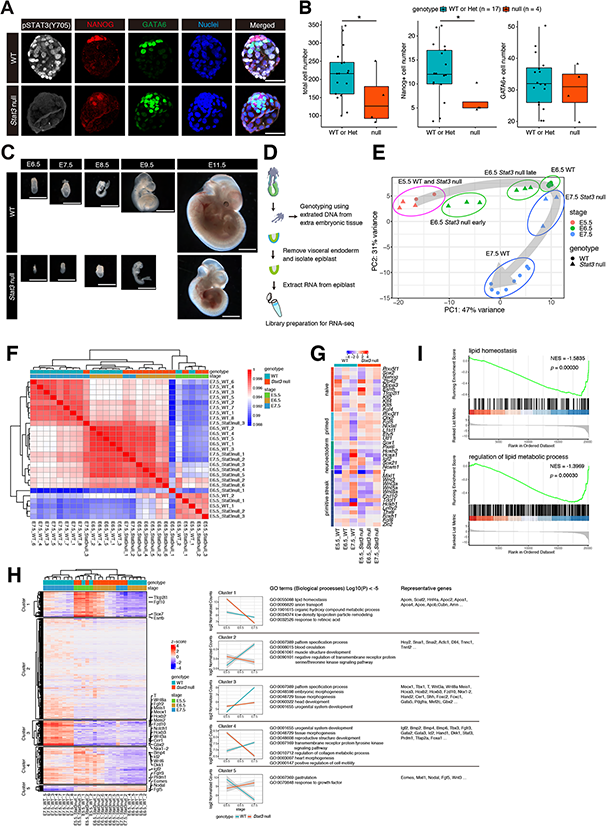
Developmental retardation of *Stat3* null embryos. (A) Immunofluorescence for pSTAT3(Y705), NANOG, and GATA6 in E4.5 WT and *Stat3* null embryos. Scale bar = 50 μm. (B) Quantification of NANOG+ and GATA6+ cells in (A) WT: n = 17, *Stat3* null: n = 4, * *P* < 0.01. (C) Morphology of *Stat3* WT and null embryos from E6.5 to E11.5. Scale bar = 1 mm. (D) Schematic of bulk RNA-sequencing from isolated epiblast cells from postimplantation embryos. (E) PCA plot of WT and *Stat3* null epiblast cells from E5.5 to E7.5. (F) Heatmap comparison and unsupervised hierarchical clustering of WT and *Stat3* null epiblast cells from E5.5 to E7.5. (G) Heatmap comparison for naïve, primed, neuroectoderm, and primitive streak genes. (H) Heatmap clustering of 1074 genes significantly different between WT and *Stat3* null epiblast (absolute log2 fold change > 2, adjusted p < 0.05). Expression pattern of each gene cluster and enriched GO biological process (p < 0.005). Representative genes are listed. (I) GSEA for lipid homeostasis and regulation of lipid metabolic process in E7.5 WT and *Stat3* null epiblast cells.

### STAT3 null epiblasts follow a normal developmental trajectory

Epiblasts were dissected from *Stat3* het *inter-se* mating at E5.5, E6.5 and E7.5, thus spanning epiblast priming and onset of gastrulation. Each was genotyped using extraembryonic tissue. Bulk RNA-sequencing was performed on denuded epiblasts to compare transcription profiles between *Stat3* WT and null embryos (Fig. 3D). While E5.5 WT and *Stat3* null epiblast cells co-localised, E6.5 null epiblasts were intermediate between WT E5.5 and E6.5; E7.5 null epiblasts grouped more closely with WT E6.5 than E7.5 epiblasts (Fig. 3E). Unsupervised hierarchical clustering confirmed that E7.5 *Stat3* null epiblasts related more closely to WT E6.5 than E7.5 (Fig. 3F). *Stat3* null epiblasts showed downregulation of naïve and upregulation of primed pluripotency genes at E6.5 (Fig. 3G), but failed to induce neuroectoderm significantly and displayed reduced intensity of selected primitive streak genes at E7.5. Differential expression analysis identified 427 upregulated and 795 downregulated genes in *Stat3* null epiblasts relative to WT (Fig. 3H). Unsupervised hierarchical clustering classified them into groups indicating characteristics of expression patterns between WT and *Stat3* null epiblast cells. Gene ontology (GO) analyses were performed for these clusters. GO terms for pattern specification and somitogenesis, relating to gastrulation and further organogenesis, were enriched in cluster 3 and highlighted delayed upregulation in *Stat3* null epiblasts over these stages (Fig. 3H). Notably, GO terms for lipid homeostasis and lipid modification were top-ranked in cluster 1. Cluster 1 genes were downregulated in *Stat3* null epiblasts at E7.5 but maintained in WT from E5.5 to E7.5. Gene set enrichment analysis (GSEA) highlighted variations in lipid homeostasis and lipid metabolism in *Stat3* null epiblast cells at E7.5 (Fig. 3I).

To assess early events causing developmental delay, we compared E5.5 WT and E6.5 *Stat3* null epiblasts. Differential expression analysis (absolute log2 fold change > 2, adjusted p < 0.05) identified 158 upregulated and 176 downregulated genes in E6.5 *Stat3* null epiblasts relative to E5.5 WT (Fig. S2A). Heatmap demonstrated that overall expression patterns of E6.5 *Stat3* null epiblasts for naïve, primed, neuroectoderm, and primitive streak genes were comparable to WT epiblasts from the previous day (Fig. S2B). GO enrichment analysis revealed that lipid localisation and fatty acid biosynthetic processes were enriched in E6.5 *Stat3* null relative to E5.5 WT (Fig. S2C). These results suggest that, while *Stat3* null epiblast cells progress along a near normal differentiation trajectory, *Stat3* deletion compromises lipid metabolism as early as E6.5.

### Primed epiblasts specifically phosphorylate S727, but not Y705 STAT3

Transcriptional Regulatory Relationships Unlabelled by Sentence-based Text mining (TRRUST) identified *Wnt* target and mesodermal regulated genes affected by *Stat3* deletion in E6.5 and E7.5 epiblasts. STAT3-controlled genes became enriched at E7.5 (Fig. 4A). Little difference was observed between WT and *Stat3* null epiblasts at E5.5, but gene expression began to diverge at E6.5, particularly when comparing WT with epiblasts from the smaller, more retarded *Stat3* null embryos (Fig. 4B). By E7.5, WT and *Stat3* null embryos were easily distinguished by morphology prior to genotyping, as shown in Fig. 3C, and this developmental divergence was reflected in the RNA-sequencing analysis (Fig. 4B). Expression of the STAT3 target Suppressor of Cytokine Signalling 3 (*Socs3*) remained low in *Stat3* null epiblasts, but also decreased in WT epiblasts at E6.5, increasing again at E7.5 (Fig. 4C). Consistent with this, pY705 was barely detectable in WT embryos soon after implantation, returning in the node region of the primitive streak at mid-late gastrulation (Fig. 4D). In contrast, pS727 was abundant throughout the epiblast from E4.5 to E7.5 (Fig. 4E). We assessed the response of WT ESCs to inhibition of signalling pathways implicated in STAT3 signalling. MEK inhibition by PD0325901 did not impair S727 phosphorylation in serum-free medium (Fig. 4F,G), whereas S727 phosphorylation is reported to depend on ERK signal in serum-containing medium (Huang et al., 2014). However, JAK inhibition only partially dampened pS727, indicating S727 can be phosphorylated by factors distinct or downstream from JAK signalling or LIF-gp130 receptors (Fig. 4F,G).

**Figure 4.**
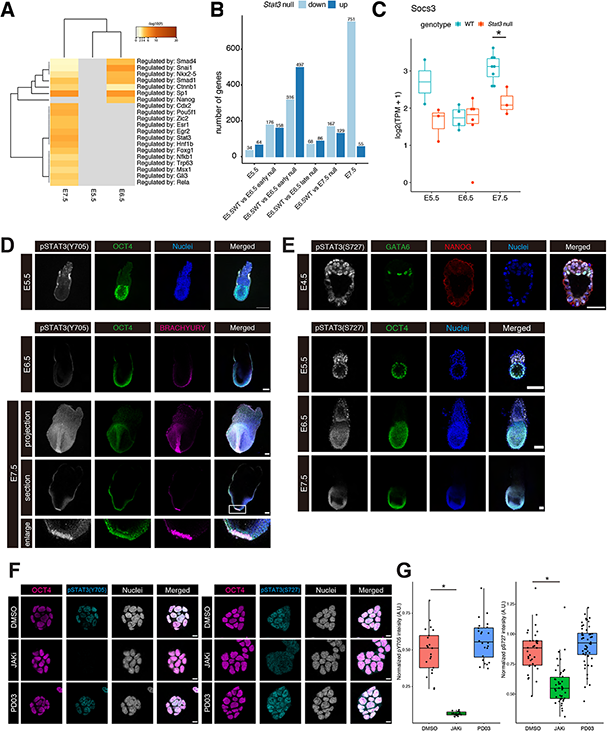
Y705 STAT3 is not phosphorylated in the post-implantation epiblast, whereas persistent phosphorylation of S727 STAT3. (A) TRRUST for differentially expressed genes in Fig.3H. (B) Numbers of up- and down-regulated genes in *Stat3* null epiblast cells (absolute log2 fold change > 2, adjusted *p* < 0.05) between E5.5 and E7.5. Note while 316 and 497 genes were identified as up- and down-regulated in E6.5 *Stat3* null early epiblast cells relative to E6.5 WT, small numbers of genes were differentially expressed in E6.5 *Stat3* null late epiblast cells. (C) log2 normalized expression levels of *Socs3* in WT and *Stat3* null epiblast cells from E5.5 to E7.5. \**P* < 0.01. (D) Immunofluorescence for pSTAT3(Y705) and OCT4 in WT E5.5 embryos and pSTAT3(Y705), OCT4, and BRACHYURY in WT E6.5 and E7.5 embryos. Scale bar = 100 μm. (E) Immunofluorescence for pSTAT3(S727), GATA6, and NANOG in WT E4.5 embryos and pSTAT3(S727) and OCT4 in WT E5.5, E6.5, and E7.5 embryos. Scale bar = 25 μm for E4.5 and 100 μm for E5.5, E6.5, and E7.5 embryos. (F) Immunofluorescence for pY705 and pS727 STAT3 in mouse ESCs. ESCs were cultured in N2B27+2i/L for 24 h, then medium exchanged to N2B27 with DMSO, JAK inhibitor, or PD03. Scale bar = 10 μm. (G) Quantification of pY705 and pS727 STAT3 intensity from immunofluorescence in (F). * *P* < 0.01. A.U., arbitrary unit.

### Self-renewing stem cell lines can be derived from *Stat3* null postimplantation epiblasts

Despite their reduced size compared with WT littermates, epiblasts from *Stat3* null embryos at E6.5 and E7.5 could be captured and propagated as EpiSC lines with high efficiency (Table S1). Their morphology in culture appeared indistinguishable from WT (Fig. 5A) and growth kinetics were similar (Fig. 5B). Whilst post-implantation epiblast cells have abundant pS727 STAT3 (Fig. 4E), neither pY705 nor pS727 was detected in WT EpiSC lines in AFX (Fig. 5C). LIF supplementation is not required for EpiSC culture (Brons et al., 2007; Tesar et al., 2007), but pY705 and pS727 were both induced by administering LIF to WT EpiSC lines (Fig. 5C). This result suggests that while FGF-ERK pathway phosphorylates S727 STAT3 in ESCs in serum/LIF medium (Huang et al., 2014), pS727 is not induced significantly by FGF in standard EpiSC culture (Fig. 5C,D). Transcriptional profiles of *Stat3* WT and null EpiSCs exhibited close clustering by means of bulk RNA-sequencing (Fig. 5E), implicating a STAT3-independent *in vitro* adaptation to self-renewal during acquisition of the mid-gastrulation anterior streak identity reported for EpiSCs, regardless of stage of origin (Kojima et al., 2014). Variance between EpiSCs and ESCs was 91%, whereas that between WT ESCs in 2i/LIF and *Stat3* null or WT ESCs without LIF was 4%, consistent with previous observations (Martello et al., 2013) (Fig. 5E). Differential expression analysis identified 39 upregulated and 99 downregulated genes in *Stat3* null EpiSCs related to WT (Fig. 5F). In concordance with a small transcriptional difference by *Stat3* deletion, heatmap for lineage markers showed a comparable expression pattern between WT and *Stat3* null EpiSCs (Fig. 5G). We conclude that STAT3 signalling is dispensable for maintenance of primed pluripotent cells *in vitro*.

**Figure 5.**
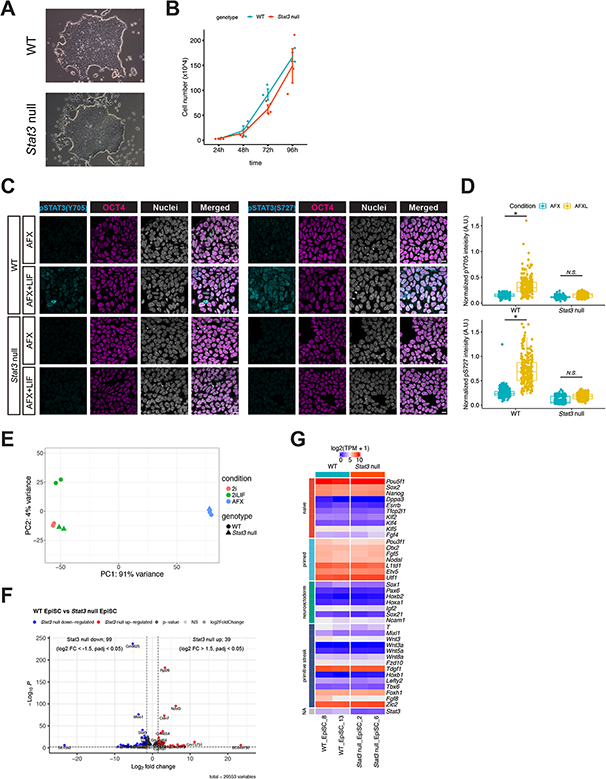
Derivation and analysis of *Stat3* WT and null EpiSC. (A) Representative bright field images of WT and *Stat3* null EpiSCs, derived from epiblast of E6.5 and E7.5 embryos (Table S1). (B) Proliferation of WT and *Stat3* null EpiSCs. (C) Immunofluorescence for pSTAT3(Y705 and S727) and OCT4 in WT EpiSC cultured in N2B27+AFX with or without LIF. Scale bar = 10 μm. (D) Quantification of pSTAT3 intensity from immunofluorescence in (C). * P < 0.01. A.U., arbitrary unit. (E) PCA plot of WT and *Stat3* null ESCs data obtained from (Betto et al., 2021) and EpiSCs from bulk RNA-seq. (F) Volcano plot of differentially expressed genes (absolute log2 fold change > 1.5, *p* < 0.05) in *Stat3* null EpiSC related to WT. (G) Heatmap comparison for naïve, primed, neuroectoderm, and primitive streak genes. Log2 (TPM + 1) was used for heatmap.

### *Stat3* null ESC aggregates exhibit defective elongation in gastruloid differentiation assays and are depleted in the differentiating region when mixed with WT ESCs

Gastruloids are organoids generated from ESCs that mimic mammalian gastrulation (van den Brink et al., 2014). *Stat3* null ESCs exhibit defects in elongation when aggregated and cultured as gastruloids, although they initiate a BRACHYURY (T)-positive pole, consistent with symmetry breaking and primitive streak instigation (Fig. 6A,B,C). As pS727 STAT3 is required for neural differentiation from ESCs (Huang et al., 2014), we generated neural tube-like structures by supplementing the culture with extracellular matrix (Veenvliet et al., 2020). By adding Geltrex between days 4 and 5 of the gastruloid protocol, WT aggregates formed neural tube-like structures, confirmed by accumulation of F-ACTIN in the SOX2-positive posterior region (Fig. 6D). *Stat3* null gastruloids elongated partially but failed to maintain BRACHYURY in the posterior region or accumulate F-ACTIN (detected by Phalloidin; Fig. 6D). Consistent with our finding that pS727 STAT3, but not pY705, was enriched in postimplantation stages, pS727 STAT3 was relatively abundant during WT gastruloid differentiation (Fig. 6E). pY705 STAT3 increased slightly at later stages, but it did not specifically correlate with BRACHYURY, suggesting that pY705 STAT3 may depend on an external signal that would normally be communicated by extraembryonic tissue *in vivo* (Fig. 6E,F,G).

**Figure 6.**
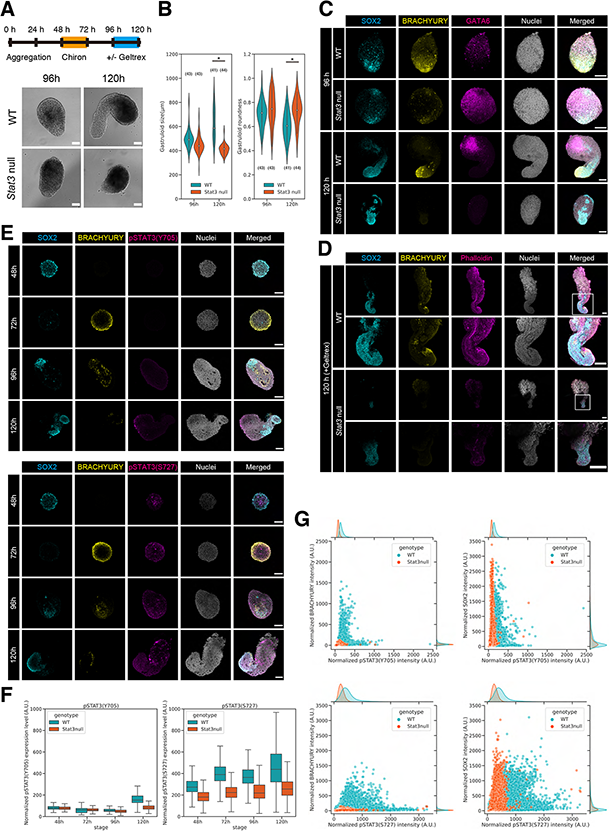
*Stat3* null gastruloids fail to elongate the posterior axis. (A) Schematic of gastruloid differentiation protocol and representative bright field image of gastruloids generated from WT and *Stat3* null ESCs at 96 h and 120 h. Scale bar = 500 μm. (B) Quantification of WT and *Stat3* null gastruloids in size (left) and shape (right) at 96 h and 120 h. Maximum feret diameter and roundness were measured from bright field images of individual gastruloids by Fiji. * *p* < 0.01. (C, D) Immunofluorescence for SOX2, BRACHYURY and GATA6 or Phalloidin in WT and *Stat3* null gastruloids at 96 h and 120 h (C), and 120 h gastruloids with neural tube-like structure (D). Scale bar = 100 μm. (E) Immunofluorescence for SOX2, BRACHYURY, pSTAT3(Y705) (top panel), or SOX2, BRACHYURY, pSTAT3(S727) (bottom panel) in WT gastruloids from 48 h to 120 h. Scale bar = 100 μm. (F,G) Normalized intensity for pSTAT3(Y705) and pSTAT3(S727) in WT and *Stat3* null gastruloids from 48 h to 120 h (F), and scatter plot for SOX2 and pSTAT3(Y705) or pSTAT3(S727), and BRACHYURY and pSTAT3(Y705) or pSTAT3(S727) in WT and *Stat3* null gastruloids at 120 h (G). pSTAT3(Y705) staining for WT; n = 3, 3, 4, 4. pSTAT3(Y705) staining for *Stat3* null; n = 4, 4, 4, 4. pSTAT3(S727) staining for WT; n = 4, 3, 4, 4. pSTAT3(S727) staining for *Stat3* null; n = 5, 4, 4, 4 for 48 h, 72 h, 96 h, and 120 h, respectively. A.U., arbitrary unit.

To begin to address the cause of the *Stat3* null developmental delay phenotype we enquired whether failure of *Stat3* null gastruloids to elongate was cell autonomous or caused by loss of STAT3 signalling. We therefore generated chimaeric gastruloids by mixing *Stat3* null with WT ESCs in various proportions. *Stat3* null cells became preferentially and progressively excluded from the BRACHYURY-positive region and were thus underrepresented in the posterior end of gastruloids (Fig. 7A,B). IF for active Caspase3 failed to distinguish significantly between WT and *Stat3* null cells during gastruloid elongation, suggesting that apoptosis was not primarily responsible for exclusion of *Stat3* null cells from the posterior region (Fig. 7B-F). While BRACHYURY was induced in *Stat3* null cells in gastruloids at 72 h, they failed to maintain its expression during subsequent culture (Fig. 7B,G,H). Furthermore, active Caspase3 was notably absent from that region in chimaeric gastruloids at 96 h, providing further support that apoptosis is not the primary cause for exclusion of *Stat3* null cells from the elongating region in the presence of differentiating WT cells (Fig. 7B-F).

**Figure 7.**
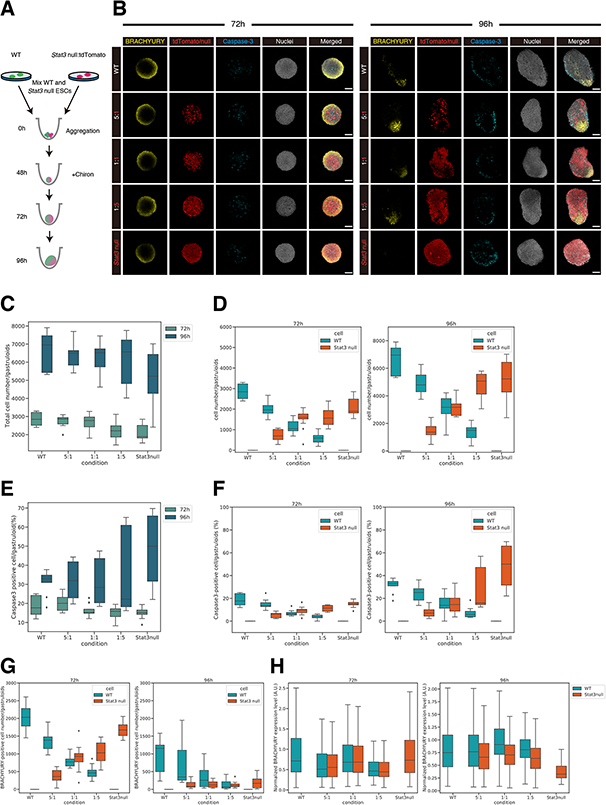
*Stat3* null cells excluded from the posterior region of gastruloids when mixed with WT cells. (A) Schematic of chimeric gastruloid differentiation using WT and tdTomato-expressing *Stat3* null ESCs. (B) Immunofluorescence for BRACHYURY, cleaved Caspase-3, and tdTomato in different ratio of chimeric gastruloids at 72 h and 96 h. Scale bar = 100 μm. (C, D) Numbers of the total cells in chimeric gastruloids (C), and numbers of WT and *Stat3* null cells in chimeric gastruloids at 72 h and 96 h (D). (E, F) Percentages of Caspase3 positive cells in chimeric gastruloids (E), and percentages of Caspase3 positive WT and *Stat3* null cells in chimeric gastruloids at 72 h and 96 h (F). (G) Numbers of BRACHYURY positive WT and *Stat3* null cells in chimeric gastruloids at 72 h and 96 h. (H) Normalized fluorescence intensity of BRACHYURY in WT and *Stat3* null cells in chimeric gastruloids at 72 h and 96 h. n = 11, 12, 13, 14, 14 at 72 h and n = 9, 12, 12, 9, 10 at 96 h for WT, 5:1, 1:1, 5:1, *Stat3* null, respectively. A.U., arbitrary unit.

## Discussion

Derivation of *Stat3* null ESCs previously facilitated interrogation of STAT3 targets and highlighted TFCP2L1 as the most significant *in vitro* (Martello et al., 2013; Ying et al., 2008). To investigate the potential role of STAT3 and TFCP2L1 in maintenance of naïve pluripotency *in vivo* we induced embryonic diapause. In contrast to the phenotype observed following deletion of LIFR or its co-receptor, gp130, which resulted in gradual loss of epiblast, but not PrE (Nichols et al., 2001), embryos lacking either STAT3 or TFCP2L1 lost virtually the entire ICM within only a few days (Fig. 2). This more dramatic phenotype may be a consequence of the precipitous depletion of epiblast, the source of PrE-inducing FGF4 (Yamanaka et al., 2010), compared with deletion of LIF receptor complex components (Nichols et al., 2001). STAT3 also promotes anti-apoptotic activity (Hirano et al., 2000), which could contribute to the enhanced PrE population reported in blastocysts supplemented with IL6 (Anderson et al., 2017; Morgani and Brickman, 2015). We conclude that STAT3 signalling is essential to maintain naïve pluripotency *in vivo* (Fig. 8) and operates as a signal transducer for pathways in addition to that induced by IL6 family cytokines.

**Figure 8.**
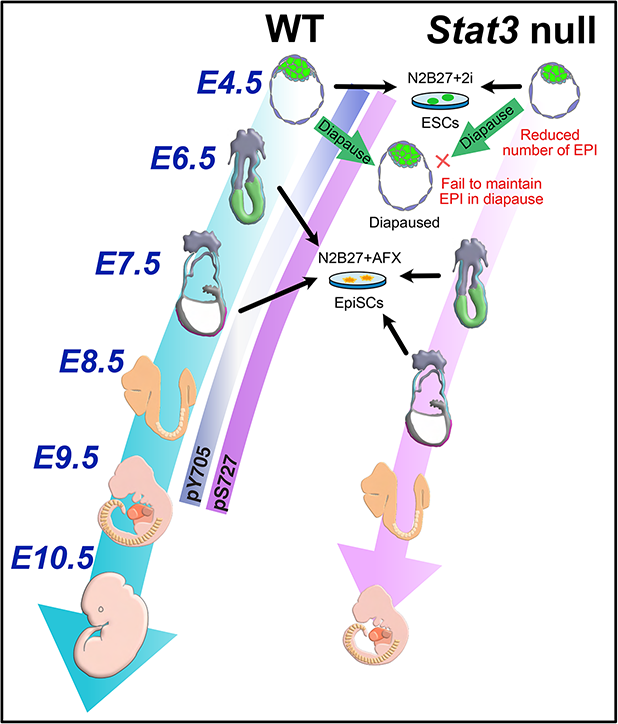
Graphical summary of *Stat3* null embryo phenotypes. In accordance with the previous report, preimplantation development of CD1 outbred *Stat3* zygotic null embryos was grossly normal, while significant reduction of epiblast and trophectoderm cell numbers was evident. When diapause was induced, *Stat3* null embryos failed to maintain NANOG+ epiblast, suggesting STAT3 signal is required for maintenance of pluripotency in diapause. After implantation, *Stat3* null embryos consistently lag behind their littermates by around one day, both morphologically and transcriptionally. pSTAT3(Y705) was undetectable in postimplantation epiblast cells until formation of the primitive streak, whereas pSTAT3(S727) was widely detected from peri- to postimplantation. EpiSCs can be derived from *Stat3* null postimplantation epiblast and maintained in N2B27 supplemented with AFX. As both phospho-variants of STAT3 were not expressed in WT EpiSCs, STAT3 signal appears dispensable for maintenance of primed pluripotency *in vitro*.

Interestingly, we recently found precocious expression of PrE markers such as *Pdgrfa, Sox17* and *Gata4* in *Stat3* null embryos at the mid blastocyst stage (E3.5), whereas emerging epiblast cells at E3.75 prematurely activated postimplantation epiblast genes *Utf1, Otx2* and its targets *Dnmt3a and b* (Betto et al., 2021), which presumably instigated reduction of FGF4 secretion. Before the time of implantation, we noted a significant decrease in the number of epiblast cells in *Stat3* null (non-diapaused) embryos compared to controls, but not the number of PrE cells (Fig. 1), potentially reflecting enhanced contribution of mutant ICM cells to this lineage, as predicted from our previous E3.5 single cell RNA sequencing data (Betto et al., 2021). However, previous observations of PrE persistence during diapause when LIFR or gp130 are deleted (Nichols et al., 2001) imply independence of this branch of STAT3 signalling for PrE maintenance. The present data implicate STAT3 signalling, via TFCP2L1, in PrE maintenance *in vivo*. Our failure to derive ESCs from *Tfcp2l1* null embryos using the strategy that proved highly successful for generation of *Stat3* null ESCs (Table S1) implies an unexpected STAT3-independent role for TFCP2L1 in transition towards *in vitro* self-renewal of pluripotent stem cells. TFCP2L1 plays a role in upregulation of *Nanog* (Ye et al., 2013), which is also indispensable for derivation of ESCs from mouse embryos (Silva et al., 2009). Interestingly, both *Nanog* and *Tfcp2l1* can be deleted from established ESCs cultured in 2i/LIF (Chambers et al., 2007; Martello et al., 2013; Yan et al., 2021), implicating compensation by the robust and redundant network of pluripotency factors assembled in ESCs *in vitro* (Dunn et al., 2014). This pathway connection may explain the failure of embryos lacking TFCP2L1 to yield ESCs and further highlights the distinct requirements for self-renewal of naïve pluripotent cells *in vivo* compared with established cell lines *in vitro*.

We observed a developmental delay in zygotically-deleted *Stat3* mutant embryos of approximately one day, consistent with the reported role for STAT3 in size regulation in later gestation and post-natal growth (Gutierrez, 2020; Shen et al., 2004). *Stat3* null embryos were otherwise apparently normal, suggesting that transition of epiblast from naïve to primed pluripotency (Nichols and Smith, 2009) and subsequent patterning of the foetus can progress, independent of STAT3 signalling (Fig. 3; Fig. 8). Consistent with this, propagation of EpiSCs derived from postimplantation epiblast does not require LIF supplementation (Brons et al., 2007; Tesar et al., 2007), but forced activation of STAT3 or administration of LIF can redirect EpiSCs to an ESC-like state (Bao et al., 2009; Yang et al., 2010). Although we could derive EpiSCs efficiently from *Stat3* null embryos, apparently indistinguishable from WT controls (Fig. 5), a role for STAT3 in differentiation emerged in gastruloids. Those generated from *Stat3* null ESCs were able to break symmetry, but failed to elongate (Fig. 6). In chimaeric gastruloids *Stat3* null ESCs were excluded from the developing ‘primitive streak’ region when mixed with WT ESCs (Fig. 7), providing further evidence that STAT3 is not required for propagation of the primed pluripotent state *in vitro*, but becomes advantageous once cells commence differentiation, as occurs in primitive streak and neural lineages. Interestingly, we observed no significant increase in apoptosis of *Stat3* null cells in chimaeric gastruloids. We therefore propose that the developmental delay phenotype revealed in embryos lacking STAT3 is a response to depletion of paracrine signalling, which may normally be provided by the overlying visceral endoderm, rather than an autocrine defect recognised by neighbouring wild type cells, leading them to instruct apoptosis, as described previously (Sancho et al., 2013).

We revealed that pY705 is negligible in the embryo between implantation and organogenesis, while persistent phosphorylation was observed at S727 (Fig. 4). This is consistent with a previous report showing requirement for pS727 in exit from pluripotency *in vitro* (Huang et al., 2014). pS727 is considered to be the mitochondrially-localised form of STAT3 (Carbognin et al., 2016; Yang and Rincon, 2016), which may suggest that the observed reduced growth of *Stat3* null embryos results from diminution of mitochondrial function. Bulk RNA-sequencing comparing epiblasts from *Stat3* null and WT embryos between time points (for example, E7.5 null with E6.5 WT) identified candidate pathways consistent with this hypothesis (Fig. 3). Lipid metabolism-related gene expression was decreased as early as E6.5 in *Stat3* null epiblast cells. Lipid metabolism is an important metabolic pathway in gastrulating embryos, and impairing nutrient uptake, such as by deletion of *Cubilin*, causes developmental arrest at gastrulation (Perea-Gomez et al., 2019). Future work will address the role of lipid metabolism in gastrulation and may provide a link to the developmental delay phenotype observed in *Stat3* null embryos at the time of gastrulation (Fig. 8).

## Materials and Methods

### Mice, husbandry and embryos

Experiments were performed in accordance with EU guidelines for the care and use of laboratory animals and under the authority of appropriate UK governmental legislation. Use of animals in this project was approved by the Animal Welfare and Ethical Review Body for the University of Cambridge and relevant Home Office licences are in place.

Mice were maintained on a lighting regime of 12:12 hours light:dark with food and water supplied *ad libitum. Stat3* mice heterozygous for replacement of exons 20-22 with Neomycin resistance (Takeda et al., 1997) were backcrossed to CD1 mice. *Tfcp2l1* heterozygous mice were generated from ESCs targeted using CRISPR strategy in E14 ESCs obtained from Jackson Labs Knockout Mouse Project (KOMP), via injection into C57BL/6 blastocysts to generate chimaeras. Male chimaeras were mated with CD1 females, grey pups were genotyped by PCR of ear biopsies and robust males selected for further backcrossing to CD1 females. Both STAT3 and TFCP2L1 mouse lines were maintained by backcrossing to CD1. Embryos were generated from *Stat3^+/-^* or *Tfcp2l1^+/-^ inter se* natural mating. Detection of a copulation plug in the morning after mating indicated embryonic day (E)0.5. Embryos were isolated in M2 medium (Sigma).

#### Genotyping

Mice were genotyped by PCR using ear biopsies collected within 4 weeks of birth and genomic DNA was extracted using Extract-N-Amp tissue prep kit (Sigma-Aldrich). Embryos were genotyped using either immune-reactivity to antibody raised against either STAT3 pY705, pS727 or TFCP2L1 in the case of those imaged for confocal analysis, or PCR analysis of trophectoderm lysate for ESC derivation or surplus extraembryonic tissue for postimplantation embryos. Amplification was carried out on around 5 μL of lysate for 35 cycles (following 95°C hot start for 10 minutes) of 94°C, 15 seconds; 60°C, 12 seconds; 72°C, 60 seconds, with a final extension at 72°C for 10 minutes. Reaction products were resolved by agarose gel electrophoresis. Primers used for genotyping PCR are listed in Supplementary Table.S4.

#### Induction of diapause

For determining the requirement for STAT3 or TFCP2L1 during maintenance of the embryo during delayed implantation, het females mated by het males were surgically ovariectomised as described previously (Nichols et al., 2001) before the embryos reached E2.5. Diapause embryos were flushed 4 days later and fixed for immunohistochemistry.

#### Derivation and culture of ESC and EpiSC

Morulae were collected from het females 2.5 days after mating by het males and used for ESC derivation as described previously (Ying et al., 2008) by culture to the blastocyst stage in KSOM supplemented with 2i, consisting of 1 μM PD0325901 and 3 μM CHIR99021, transfer of ICMs isolated by immunosurgery to 48-well plates containing 2i in N2B27 medium, one per well. WT and *Stat3* null ESCs were expanded and maintained in N2B27 supplemented with 2i or 2i/LIF on gelatin-coated plates at 37°C in 7% CO_2_ and passaged by enzymatic disaggregation every 2-3 days. To examine phosphorylation of STAT3 in ESCs, WT cells were cultured in N2B27 with 2i/LIF for 24 h then N2B27 supplemented with DMSO, 1 μM PD0325901, or 0.5 μM JAK inhibitor I (JAKi, Calbiochem) for 2 h.

For EpiSC derivation, epiblasts were manually isolated from E6.5 or E7.5 embryos obtained from *Stat3* het intercross by removal of extraembryonic ectoderm and visceral endoderm by means of flame-pulled Pasteur pipettes of appropriate diameter. Genotyping was performed using genomic DNA extracted from extraembryonic tissue as described above. Isolated epiblasts were plated on fibronectin-coated plates in N2B27 medium supplemented with AFX, consisting of 12 ng/mL FGF2, 20 ng/mL Activin A and 10 μM XAV929. EpiSCs were maintained in N2B27 supplemented with AFX in fibronectin coated-plates at 37°C in 5% O_2_ and passaged every 3-4 days. To examine phosphorylation of STAT3 in EpiSCs, WT and *Stat3* null EpiSCs were cultured with 10 ng/mL LIF in N2B27 +AFX medium for 1 h.

#### Transfection

To generate stable tdTomato-expressing WT or *Stat3* null ESC lines, cells were plated in 24-well plates the day before transfection. 2 μg of plasmid DNA and 5 μL of Lipofectamine2000 (Thermo Fischer Scientific #11668019) were incubated in 100 μL N2B27 for 20 min and then mixed and incubated for 20 min at room temperature. The mixture was added to cells and cultured for 3 hr, then exchanged for fresh medium. After 24 h, cells were passaged to gelatin-coated 6-well plates and puromycin (1 μg/mL) selection performed. Expanded colonies were picked manually and cell lines uniformly expressing tdTomato selected by flow cytometry.

#### Generation of gastruloids

Gastruloids were generated as previously reported (Baillie-Johnson et al., 2015) with some modifications. Briefly ESCs were cultured in N2B27 with 2i or 2i/LIF on gelatin-coated plates and dissociated into single cells by Accutase. They were then centrifuged at 1,300 rpm for 3 min and washed twice with PBS. Cells were resuspended with pre-warmed N2B27 and cell numbers counted using a hematocytometer. To make aggregates, 200-300 cells in 40 μL N2B27 were transferred to U-bottom 96-well plate using multichannel pipette and cultured for 48 h at 37°C in 5% CO_2_. At 48 h of culture, secondary medium was prepared by adding 3 μM CHIR99021 into 37°C and 5% O_2_ pre-equilibrated N2B27 and 150 μL added to each well using multichannel pipette. At 72 h of culture, the medium was exchanged for pre-equilibrated N2B27 and repeated until 120 h. To make neural tube-like structures, gastruloids were embedded in 2% Geltrex between 96 h and 120 h (Veenvliet et al., 2020). To make chimaeric gastruloids, WT and *Stat3* null ESCs were mixed in specific ratios and aggregated in 96-well plate.

#### Immunofluorescence

Embryos were fixed in 4% paraformaldehyde (PFA) for 30 min at room temperature (RT), followed by washing in 0.5% PVP/PBS. Embryos were permeabilized in 0.5% Triton X-100 in PBS for 15 min and blocked with 2% donkey serum, 2.5% BSA, and 0.1% Tween20 in PBS for 1 h at RT. For phosphorylated-STAT3 staining, permeabilization was performed in absolute methanol for 10 min at −20°C. Primary antibodies were diluted in blocking solution and incubated overnight at 4°C. After washing embryos in 0.1% Tween 20 in PBS, embryos were incubated with Alexa Fluor-conjugated secondary antibodies (Thermo) for 1 h at RT. Nuclear staining was carried with Hoechst33342 (Thermo). Primary and secondary antibodies used are listed on Table S3.

For IF of ESCs and EpiSCs, cells were cultured on fibronectin coated ibidi-treated 8-well chamber slide (ibidi). Cells were fixed in 4% PFA for 15 min at RT, followed by washing in PBS. Cells were permeabilized in absolute methanol for 10 min at −20°C and then washed in PBS. Blocking reaction was carried out with 2% donkey serum, 2.5% BSA, and 0.1% Tween20 in PBS for 1 h at RT. Primary antibodies were diluted in blocking solution and incubated overnight at 4°C. After washing cells in 0.1% Tween 20 in PBS, cells were incubated with Alexa Fluor-conjugated secondary antibodies (Thermo) for 1 h at RT. Nuclear staining was carried with Hoechst33342 (Thermo).

Gastruloids for IF were collected from each stage and fixed in 4% PFA for 30 min at RT, followed by washing in 0.5% PVP/PBS. Gastruloids were then taken through the same IF procedure as used for embryos.

#### Imaging and image analysis for embryos

Images for embryos were acquired using TCS SP5 (Leica) confocal microscope and processed with ImageJ. Quantification of immunofluorescence images of preimplantation and diapause embryos was achieved using a computational pipeline to extract data on the intensity of antibody staining in individual nuclei. A 3D StarDist segmentation model was trained with a manually-corrected modular interactive nuclear segmentation (MINS) annotated dataset of E3.5-E4.5 mouse blastocysts (Lou et al., 2014; Schmidt et al., 2018; von Chamier et al., 2021). Nuclear segmentation of embryos, generated by applying the pre-trained model, was used as a mask to measure fluorescence intensity, xyz position, and morphological parameters (e.g., volume) of each nucleus (Ollion et al., 2013; Pietzsch et al., 2015; Schmid et al., 2010). Assessment of the histograms for signal intensity in the DAPI channel and nuclear volume allowed thresholds to be set to remove erroneously segmented nuclei. Channel-specific background subtraction was performed to account for varying levels of noise between antibodies. Violin plots were generated using the ggplot2 package in R and statistical significance between embryo groups was assessed via pair-wise unpaired student’s t-tests. Nuclei were visualised in a 2D gene expression space based on NANOG and STAT3 intensity and selected nuclei were plotted back into the ‘in silico’ embryos based on their positional information and colour coded by STAT3 intensity. ‘In silico’ embryos were generated using xyz centroid information from the segmentation mask and colour coded according to STAT3 intensity.

#### Imaging and image analysis for ESCs and EpiSCs

Images of ESCs and EpiSCs was acquired with LSM980 (Zeiss) confocal microscope and processed with ImageJ. Segmentation of nucleus and quantification of pSTAT3 levels in mouse ESCs and EpiSCs were performed using CellProfiler 4.2.1 (McQuin et al., 2018). Mean intensity value of pSTAT3 in each nuclear was normalized by mean intensity of Hoechst channel.

#### Imaging and image analysis for gastruloids

To quantify size and roundness, gastruloids were imaged using Leica DMI4000 microscope and analysed with imageJ Fiji distribution (Schindelin et al., 2012). Morphometric analysis was performed as described (De Belly et al., 2021). Briefly, images were converted to 8-bit grayscale and thresholded manually to segment gastruloid outlines. Analysis of particle function was performed and “Feret’s diameter” and “shape descriptions” were used for analysis. Fluorescence imaging of gastruloids was performed using LSM 880 (Zeiss) or Stellaris8 (Leica) confocal microscope and processed with ImageJ. Nuclear segmentation and quantification of immunofluorescence intensity of gastruloids was carried out with Imaris software (Oxford Instruments). Briefly, images were normalized using “Normalize Layer” function and then “Spot” function was used for the nuclear segmentation with estimated x-y size 6 - 8 μm. Mean Intensity value from segmented nuclear was used for the quantification.

#### RNA-sequencing

For low-input RNA-sequencing, embryos from E5.5 to E7.5 were dissected from *Stat3* heterozygous intercrosses, and extraembryonic ectoderm and visceral endoderm removed manually. Extraembryonic ectoderm was used for genotyping. RNA was isolated from single epiblasts with Picopure RNA-isolation kit (Thermo). For library construction, SMARTerR Stranded Total RNA-seq kit v2-Pico InputMammalian (Takara Clontech) was used. For bulk RNA-sequencing for EpiSCs, total RNA was isolated using RNeasy Mini Kit (QIAGEN). Ribo-Zero rRNA Removal kit (Illumina) was used for rRNA removal and NEBNext Ultra II DNA Library Prep Kit for Illumina was used for the library construction. Reads were aligned to the mouse (GRCm38/mm10) reference genome using HISAT2(v2.2.1). StringTie (v2.1.4) was used for gene counts. R environment and Bioconductor were used for all RNA-seq analysis. Normalization and differential expression analysis were carried out with DESeq2 package (v1.30.1). Transcripts with absolute log2 fold change > 2 and the p-value adjusted (padj) < 0.05 considered as differentially expressed gene. Regularized log transform (rlog) normalized values were used for principal component analysis (PCA) and heatmap. GO-term enrichment analysis and TRRUST was performed with Metascape (Zhou et al., 2019). WT and *Stat3* null ESC bulk RNA-seq data was obtained from (Betto et al., 2021).

## Acknowledgements

We wish to thank William Mansfield for generation of Tfcp2l1+/- mouse line; Kenneth Jones for lab management and help with genotyping; Thomas Burdon and Lawrence Bates for helpful discussion on the project; Shizuo Akira for providing *Stat3+/-* mice; Peter Humphreys, Darran Clement, Ann Wheeler and Laura Murphy for advanced imaging and analysis; Maike Paramor and Vicky Murray for sequencing; UBS, Cambridge for animal husbandry. This work was supported by the University of Cambridge, BBSRC project grant RG74277 and funded in part by Wellcome Trust (Grant number: 203151/Z/16/Z). T.A. is supported by JSPS Overseas Research Fellowship and UEHARA Memorial Foundation Fellowship.

For the purpose of Open Access, the author has applied a CC BY public copyright licence to any Author Accepted Manuscript version arising from this submission.

## Data Availability

Bulk RNA-seq data will be deposited in Gene Expression Omnibus.

Fiji macro and R codes will be made available at https://github.com/SKraunsoe/STAT3_embryo_segmentation_analysis.

---

**Fig. S1.**
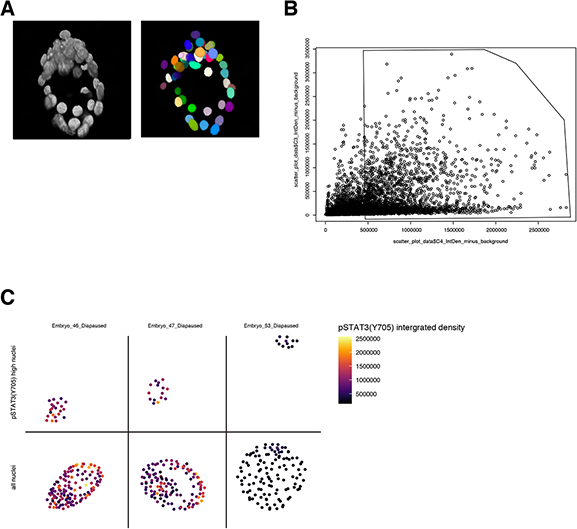
Quantification output from image analysis pipeline. (A) 3D view of the DAPI channel and segmentation output for an embryo created using the 3D viewer plugin in Fiji (Schmid et al., 2010). Videos of 3D rotations available in the supplementary figures. (B) Nuclei colour-coded by stage dispersed in a 3D co-expression space based on integrated density (sum across all pixel values in the 3D object) of Nanog and pSTAT3 and in each segmented nucleus. The polygon selection encloses cells with a pSTAT3 raw integrated density above 5 x 105. (C) Distribution of the selected high pSTAT3 cells within the polygon from (B) in their embryo of origin for 3 representative diapaused embryos (top) with all nuclei shown below.

**Fig. S2.**
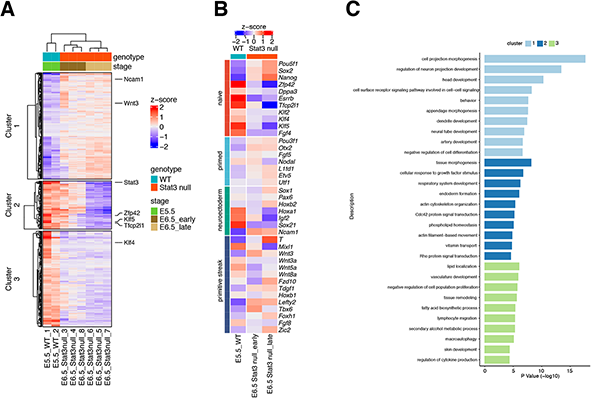
Deficiency of lipid metabolism is the earliest phenotype of *Stat3* null epibast cells at E6.5. (A) Heatmap for differentially expressed genes (absolute log2 fold change > 2, adjusted p < 0.05) between E5.5 WT and E6.5 *Stat3* null epiblast cells. (B) Heatmap comparison for naïve, primed, neuroectoderm, and primitive streak genes. (C) Enriched GO biological process (p < 0.005) for clusters in (A).

**Table S1.** Derivation of ESCs from *Stat3* and *Tfcp2l1* heterozygous intercrosses and EpiSCs from *Stat3* heterozygous intercrosses.

|  | +/+ | +/- | -/- | Total ICM |
| --- | --- | --- | --- | --- |
| ESCs | 9 | 32 | 13 | 56 |

ESCs derivation from ICM of Tfcp2l1 heterozygous intercrosses
|  | +/+ | +/- | -/- | Total ICM |
| --- | --- | --- | --- | --- |
| ESCs | 6 | 49 | 0 | 65 |

EpiSCs derivation from epiblast of Stat3 heterozygous intercrosses
| EpiSCs | +/+ | +/- | -/- | Total epi |
| --- | --- | --- | --- | --- |
| E6.5 | 3 | 11 | 3 | 17 |
| E7.5 | 3 | 7 | 3 | 13 |

**Table S2.** Numbers of *Stat3* WT, Het, and null embryos from *Stat3* heterozygous intercrosses.

|  | +/+ | +/- | -/- | empty<br>decidua |
| --- | --- | --- | --- | --- |
| E3.5 | 14 | 36 | 16 |  |
| E4.5 | 22 | 54 | 21 |  |
| E5.5 | 5 | 17 | 6 |  |
| E6.5 | 21 | 42 | 18 | 2 |
| E7.5 | 17 | 28 | 10 | 4 |
| E8.5 | 10 | 25 | 5 | 4 |
| E9.5 | 14 | 28 | 2 | 5 |
| E10.5 | 6 | 12 | 1 | 3 |
| E11.5 | 10 | 19 | 4 | 3 |

**Table S3.** List of primary and secondary antibodies.

| Antibodies | Supplier | Cat# |
| --- | --- | --- |
| Mouse monoclonal anti-Oct3/4 (C-10) | Santa Cruz | Cat#SC-5279; RRID:AB_628051 |
| Goat polyclonal anti-Brachyury | R&D systems | Cat#AF2085; RRID:AB_2200235 |
| Rat monoclonal anti-Nanog | eBioscience | Cat#14-5761-80; RRID:AB_763613 |
| Rat monoclonal anti-OCT3/4 | eBioscience | Cat#14-5841-82;RRID: AB_914301 |
| Rat monoclonal anti-SOX2 | eBioscience | Cat# 14-9811-82, RRID:AB_11219471 |
| Rabbit anti-phospho-(Tyr705) STAT3 | Cell Signaling Technology | Cat#9145; RRID:AB_2491009 |
| Rabbit anti-phospho-(Ser727) STAT3 | Cell Signaling Technology | Cat#34911; RRID:AB_2737598 |
| Goat anti-Tfcp2l1 | R&D systems | Cat# AF5726; RRID:AB_2202564 |
| Goat anti-GATA6 | R&D systems | Cat#AF1700; RRID:AB_2108901 |
| Goat anti-Brachyury | R&D systems | Cat#AF2085; RRID:AB_2200235 |
| Rabbit anti-Ki67 | Cell Signaling Technology | Cat#9129S; RRID:AB_2687446 |
| Rabbit anti-cleaved-Caspase-3 | Cell Signaling Technology | Cat#9664S; RRID:AB_2070042 |
| Mouse anti-CHOP | Cell Signaling Technology | Cat#2895; RRID:AB_2089254 |
| Donkey anti-rat Alexa488 | Thermo | Cat# A-21208, RRID:AB_2535794 |
| Donkey anti-mouse Alexa488 | Thermo | Cat# A-21202, RRID:AB_141607 |
| Donkey anti-rabbit Alexa555 | Thermo | Cat# A-31572, RRID:AB_162543 |
| Donkey anti-goat Alexa647 | Thermo | Cat# A-21447, RRID:AB_2535864 |
| Donkey anti- Mouse Alexa647 | Thermo | Cat# A-31571, RRID:AB_162542 |
| AlexaFluor 647 Phalloidin | Cell Signaling Technology | Cat#8940 |

**Table S4.**
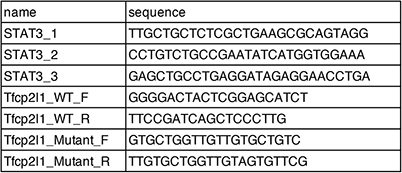
List of primers for genotyping.

